# Gastroprotective Effects of *Aleurites moluccanus* (Candlenut) Oil Against Aspirin-Induced Gastric Injury in Rats: A Dose-Dependent Histopathological Evaluation

**DOI:** 10.1101/2025.10.16.682748

**Authors:** Eman Sutrisna, Mohammad Rizky Ariesto, Hidayat Sulistyo, Iwan Purnawan

## Abstract

Nonsteroidal anti-inflammatory drugs (NSAIDs), such as aspirin, contribute significantly to drug-induced gastrointestinal complications globally, primarily through prostaglandin inhibition and oxidative stress mechanisms. The search for safer, plant-based alternatives has intensified, particularly in resource-limited settings. This study aimed to evaluate the gastroprotective potential of *Aleurites moluccanus* (candlenut) oil in a rat model of aspirin-induced gastric injury. Twenty-five male Wistar rats were randomly assigned to five groups: a healthy control, a negative control, and three treatment groups receiving *A. moluccanus* oil at doses of 0.5, 1.0, and 2.0 mL/200 g BW (equivalent to 2.5, 5.0, and 10 mL/kg BW, respectively). All groups except the healthy control were administered aspirin (600 mg/kg) to induce gastric lesions, followed by 28 days of treatment. Histopathological analysis revealed dose-dependent improvements in mucosal integrity and reduced inflammatory infiltration, with the highest dose group showing near-normal histological features. Although not all intergroup differences reached statistical significance, the consistent histological trend suggests a biologically meaningful protective effect. These findings support the therapeutic potential of *A. moluccanus* oil as a locally available, low-toxicity phytotherapeutic candidate for NSAID-induced gastritis and warrant further investigation into its molecular mechanisms and safety profile.

## INTRODUCTION

Gastritis refers to inflammation of the gastric mucosa, a condition that may present acutely or chronically and is commonly associated with epigastric pain, nausea, and indigestion (1). According to global estimates, more than half of the world’s population is affected by some form of gastritis, with a higher prevalence observed in regions with limited access to healthcare and sanitation (2). Chronic gastritis, if left untreated, may lead to serious complications such as peptic ulcer disease, gastrointestinal bleeding, or even gastric cancer (3). The economic burden of gastritis includes not only the direct costs of diagnosis and treatment but also the indirect costs associated with lost productivity and decreased quality of life (4). Given its high prevalence and potential for complications, gastritis continues to pose a major public health concern that necessitates improved preventive and therapeutic strategies(5).

NSAIDs such as aspirin are among the most widely used medications globally due to their analgesic, antipyretic, and anti-inflammatory effects(6). However, their therapeutic action is closely tied to their adverse effects on the gastric lining, primarily through the inhibition of cyclooxygenase (COX) enzymes, which are essential for the synthesis of protective prostaglandins in the stomach(6). This prostaglandin suppression reduces mucus and bicarbonate secretion, impairs mucosal blood flow, and compromises the epithelial barrier, making the stomach more vulnerable to injury(7,8). Additionally, aspirin increases gastric oxidative stress by promoting the generation of reactive oxygen species (ROS), which further exacerbate inflammation and tissue damage(9). Consequently, long-term or high-dose NSAID use is strongly associated with the development of gastritis, gastric erosions, and ulcerative lesions in susceptible individuals(10).

Proton pump inhibitors (PPIs), histamine-2 receptor antagonists (H2RAs), and antacids remain the mainstay treatments for gastritis and peptic ulcer disease, often providing symptomatic relief and mucosal healing(11). However, chronic use of these agents, particularly PPIs, has been associated with adverse effects such as nutrient malabsorption, increased risk of bone fractures, kidney injury, and alterations in gut microbiota(12). Moreover, accessibility to these drugs remains a challenge in low-resource settings where over-the-counter availability may be limited or cost-prohibitive for long-term use. There is also growing concern regarding the diminished efficacy of these therapies in certain patients due to tolerance development or inappropriate usage(11). These limitations underscore the urgent need to explore alternative therapeutic options that are not only effective but also safer, economically viable, and compatible with long-term administration.

Over the past two decades, the pharmacological potential of medicinal plants has been extensively investigated as a response to the limitations of synthetic drugs, particularly in chronic conditions such as gastritis(13)(14). These plant-based therapies offer a wide range of bioactive compounds—such as flavonoids, alkaloids, terpenoids, and phenolics—that exhibit anti-inflammatory, antioxidant, and cytoprotective activities through multi-target mechanisms(15). Unlike conventional drugs that often act on a single pathway, phytochemicals tend to modulate multiple biological pathways simultaneously, making them attractive candidates for complex diseases with multifactorial pathogenesis. Moreover, traditional medicinal systems, including Ayurveda, Traditional Chinese Medicine, and Jamu (Indonesian traditional medicine), have long utilized herbal remedies for gastrointestinal disorders, supporting their historical safety profile(14). The relatively low incidence of side effects and potential for local availability further strengthen the case for integrating natural products into modern gastroprotective treatment strategies(16).

Belonging to the Euphorbiaceae family, Aleurites moluccanus has been traditionally employed in various regions of Southeast Asia to manage ailments including digestive discomfort, skin inflammation, and joint pain(17). Phytochemical studies have identified that candlenut oil contains linoleic acid, oleic acid, flavonoids, and polyphenols—compounds known for their free radical scavenging and anti-inflammatory capabilities(18). These bioactive components may confer mucosal protection by reducing oxidative stress, modulating pro-inflammatory cytokines, and enhancing mucosal healing, mechanisms that are crucial in preventing or mitigating NSAID-induced gastric injury(19). In vivo and in vitro studies on related plant species have demonstrated similar effects, thereby encouraging further pharmacological exploration of candlenut’s therapeutic potential. Despite its long-standing use in folk medicine, the scientific validation of A. moluccanus as a gastroprotective agent remains limited, providing a strong rationale for systematic investigation.

Candlenut (Aleurites moluccanus) oil has gained attention for its reported anti-inflammatory, antimicrobial, and antioxidant activities, which are attributed to its diverse phytochemical constituents(20). Prior research has primarily focused on its dermatological and anti-inflammatory applications, including its effects on skin barrier repair, wound healing, and modulation of inflammatory mediators in vitro(21). However, despite its promising bioactivity profile, investigations into its potential benefits in gastrointestinal pathology— particularly in NSAID-induced gastric mucosal injury—are extremely limited and largely anecdotal. To date, no published in vivo studies have systematically evaluated the gastroprotective efficacy of candlenut oil in experimental models of drug-induced gastritis, leaving a critical gap in evidence. Therefore, the present study aims to assess the protective effect of A. moluccanus oil on aspirin-induced gastric mucosal injury in rats, with particular emphasis on histopathological outcomes and dose-dependent efficacy.

In many low- and middle-income countries, access to conventional gastroprotective medications such as proton pump inhibitors and H2 receptor antagonists remains constrained by economic, geographic, and systemic healthcare limitations(22,23). Moreover, patients with contraindications to standard pharmacotherapy—such as those with chronic kidney disease, electrolyte imbalance, or polypharmacy—would greatly benefit from safer and more biocompatible alternatives (24,25). Developing therapeutic agents derived from traditional medicinal plants like Aleurites moluccanus aligns with the global shift toward integrative and personalized medicine, particularly in the context of sustainable and locally sourced healthcare solutions. By validating the efficacy of candlenut oil through preclinical models, this research can lay the groundwork for future

## MATERIAL AND METHODS

### Study Design

This study employed a true experimental design with a post-test-only control group to evaluate the gastroprotective potential of *Aleurites moluccanus* (candlenut) oil in aspirin-induced gastritis in rats. A total of five experimental groups were established, consisting of a healthy control group (Group A), a negative control group receiving aspirin without treatment (Group B), and three treatment groups (Groups C, D, and E) that received candlenut oil at doses of 0.5, 1.0, and 2.0 mL/200 g BW (equivalent to 2.5, 5.0, and 10 mL/kg BW), respectively. The dose range was informed by an unpublished preliminary screening study in rats, which indicated no acute toxicity at these levels and suggested a dose-dependent gastroprotective trend. The highest dose was selected below the threshold of reported toxicity in rodents (26), while lower doses were included to enable exploration of a broad therapeutic window for subsequent optimization. Aspirin at a dose of 600 mg/200 g BW (equivalent to 3,000 mg/kg BW) was used to induce gastritis in all groups except the healthy control Rats were randomly assigned to groups using a simple randomization method generated by a computerized sequence to minimize allocation bias and ensure homogeneity.

### Animal Model and Housing Conditions

A total of 25 male Wistar rats (*Rattus norvegicus*), aged 8–10 weeks, weighing between 190 and 210 grams, were obtained from a certified breeding facility. All animals were acclimatized for 14 days prior to the experiment in a temperature-controlled room (22 ± 2°C), under a 12-hour light/dark cycle, with access to a standard pellet diet and water ad libitum. Rats were housed in standard polypropylene cages, with five animals per cage, and environmental enrichment was provided. The experimental protocol followed the 2021 Indonesian Food and Drug Authority (BPOM) guidelines on preclinical pharmacodynamic testing of traditional medicines and complied with the ARRIVE 2.0 guidelines for animal research reporting. Ethical clearance was obtained from the Medical Research Ethics Committee, Faculty of Medicine, Universitas Jenderal Soedirman (Approval No. 002/KEPK/PE/XI/2022).

The grouping and treatments are summarized in Table 1.

**Table 1.**
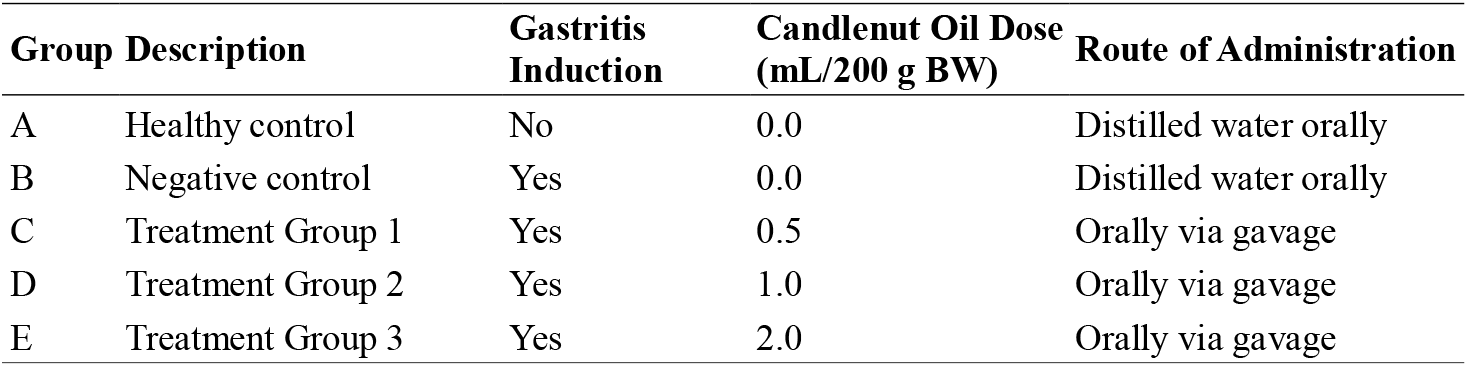
Animal grouping and treatment regimen.

**Table 2.** Modified Rogers scoring criteria for gastric mucosal damage.

| Score | Description |
| --- | --- |
| 0 | Intact mucosa with no visible lesion or inflammation |
| 1 | Mild epithelial disruption with minimal inflammatory cell infiltration |
| 2 | Moderate mucosal erosion, mild edema, and focal hemorrhage |
| 3 | Extensive epithelial loss with submucosal edema and inflammatory response |
| 4 | Severe ulceration with widespread hemorrhage and dense inflammatory infiltrate |

**Table 3.** Roger’s Score for Histopathological Feature of the Rat Gastric Mucosa.

| Intervention Groups (n=5) | Means ± SD | Min | Max | p-value* |
| --- | --- | --- | --- | --- |
| A | 2,20±0,837 | 1 | 3 | 0,001 |
| B | 5,40±1,673 | 4 | 8 | |
| C | 5,00±0,707 | 4 | 6 | |
| D | 4,20±0,837 | 3 | 5 | |
| E | 4,00±1,000 | 3 | 5 | |

\*One-way Anova Test; Post Hoc: B vs. A: $p < 0.01$ ; B vs. C, D, E: $p > 0.05$ ; E vs. Group A: $p = 0.096$

### Preparation of Candlenut Oil

Candlenut seeds (*Aleurites moluccanus*) were sourced from cultivated plantations in Tanjung Gusta, Sunggal District, Deli Serdang Regency, North Sumatra, Indonesia. Seeds were washed and sun-dried under a black cloth to prevent photodegradation. Once dried, the seeds were roasted in a convection oven at 100 °C for 60 minutes until uniformly browned. Roasted seeds were ground into an oily paste using a blender. Oil was extracted by cold pressing, followed by muslin cloth filtration, and collected in sterile amber glass bottles. The average oil yield was 27.5% w/w of dried seeds, calculated as the weight of oil obtained per weight of roasted seeds.

Roasting facilitated oil release by reducing seed moisture, whereas subsequent cold pressing was employed to minimize thermal degradation of polyunsaturated fatty acids and other bioactive compounds. Extracted oil was stored at ambient temperature in the dark and used within 7 days. This processing approach has been reported to reduce phorbol esters while maintaining the relative levels of polyunsaturated fatty acids (linoleic and oleic acids) and flavonoids, thereby preserving the biological activity of candlenut oil (27).

### Induction of Gastritis and Treatment Procedure

Following acclimatization, rats were randomly assigned into their respective groups. To induce gastric mucosal injury, aspirin was administered orally via gavage at a dose of 600 mg/200 g BW once daily for 28 consecutive days, except in the healthy control group which received distilled water. Treatment groups received candlenut oil at their assigned doses concurrently with aspirin, administered via oral gavage in a volume not exceeding 2.5 mL per 200 g BW to maintain gastric tolerability. The treatment period lasted 28 days. The treatment period lasted 28 days to simulate sub-chronic NSAID exposure, as previously reported in aspirin-induced gastritis models (28). This duration allows evaluation of cumulative mucosal injury and protective effects of interventions beyond the acute phase On day 29, all animals were lightly anesthetized with ether inhalation and euthanized via cervical dislocation. A midline laparotomy was performed to isolate the stomach. The organ was incised along the greater curvature, rinsed in saline, and immersed in 10% neutral buffered formalin (NBF) for 48 hours for fixation. Prior to histological processing, gross examination of gastric mucosa was performed to assess hyperemia and surface erosions as preliminary indicators of mucosal injury.

### Histopathological Processing and Evaluation

Fixed stomach tissues were processed using standard paraffin embedding techniques. Sections of 5 µm thickness were prepared using a Leica RM2125 microtome and stained with hematoxylin-eosin (HE). Histological analysis was performed using an Olympus BX51 light microscope under 100x and 400x magnifications. Evaluation of gastric mucosal integrity was performed by a blinded histopathologist using a semi-quantitative scoring system adapted from Rogers (2012), which categorized lesions based on epithelial disruption, edema, hemorrhage, and leukocyte infiltration. Histological evaluation was independently performed by two blinded observers, including one board-certified pathologist. Both were unaware of group allocation during scoring, thereby providing internal validation of the assessment

Three representative fields per section were scored, and the average score per animal was calculated.

### Statistical Analysis

All statistical analyses were performed using SPSS version 26.0 (IBM Corp., Armonk, NY, USA). Normality of data distribution was tested using the Shapiro-Wilk test. Homogeneity of variance was assessed using Levene’s test. If data met parametric assumptions, intergroup comparisons were conducted using one-way analysis of variance (ANOVA) followed by Tukey’s post hoc test. If non-parametric conditions were met, the Kruskal-Wallis test followed by Mann–Whitney U test was applied. Results were expressed as mean ± standard deviation (SD), and a p-value < 0.05 was considered statistically significant.

## RESULTS AND DISCUSSION

### Histopathological Changes in Gastric Mucosa

No systemic side effects were observed during the 28-day treatment period. Rats across all groups maintained stable body weights and exhibited normal grooming and locomotor activity, suggesting that candlenut oil administration did not induce overt systemic toxicity. Representative histological images of gastric mucosa from all groups are shown in **Figure 1**. The healthy control group (Group A) exhibited preserved gastric architecture with intact surface epithelium, well-defined gastric pits, and minimal inflammatory infiltration. Minor histological variations observed in this group, such as slight leukocyte infiltration, were considered within the normal physiological range and did not represent pathological lesions. In contrast, the negative control group (Group B), which received aspirin alone, displayed marked mucosal disruption characterized by epithelial erosion, obliteration of gastric pits, and dense submucosal infiltration extending into the muscularis layer, consistent with NSAID-induced gastritis.

**Figure 1.**
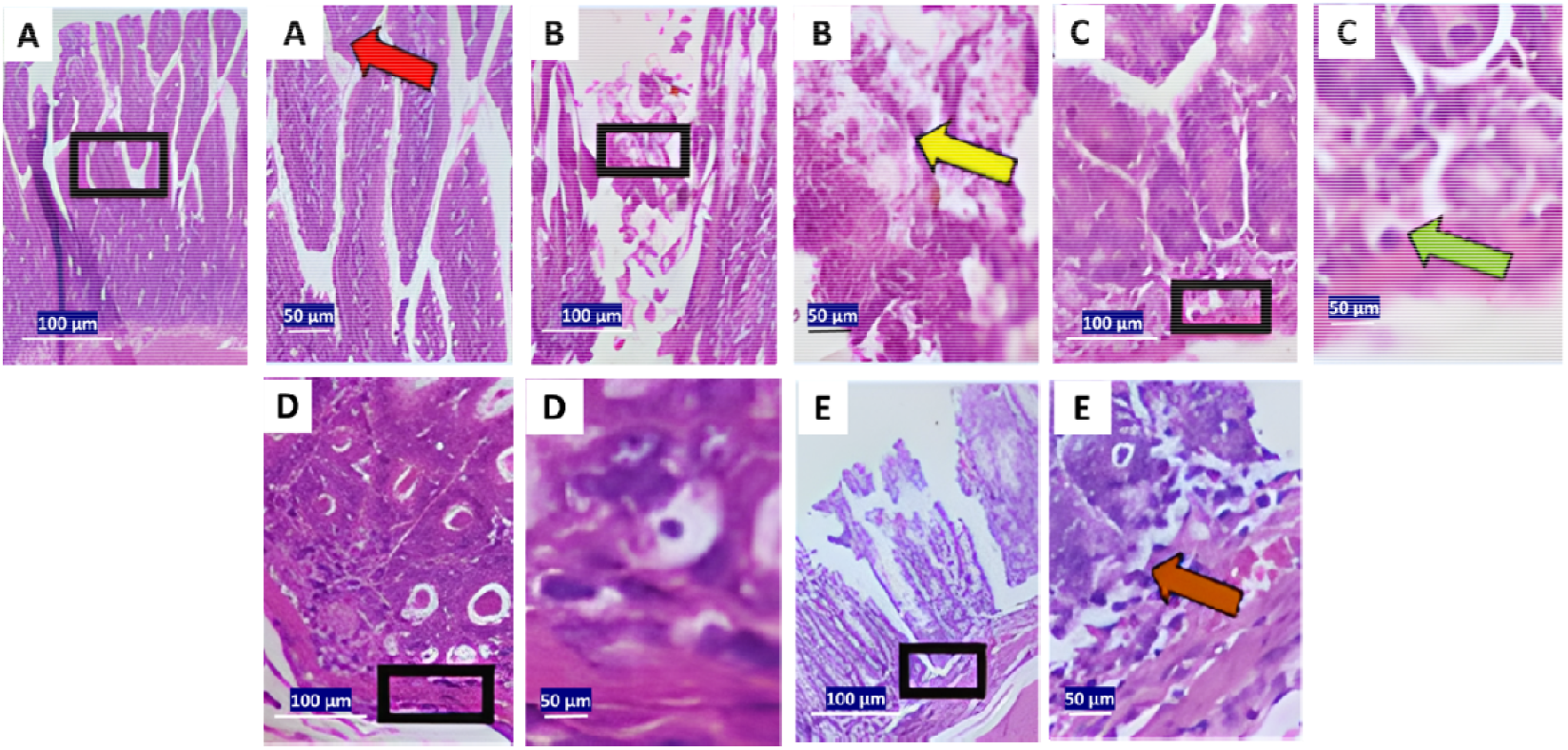
Representative hematoxylin–eosin (H&E) stained sections of rat gastric mucosa across experimental groups A–E at 100× (left) and 400× (right). Black boxes in 100× panels indicate regions of interest (ROIs) shown at higher magnification in the adjacent 400× panels. Red arrows: intact epithelium; yellow arrows: mucosal erosion; green arrows: submucosal inflammation; blue arrows: lymphoid nodules; brown arrows: transmural infiltration. Scale bars: 100 µm (100×) and 50 µm (400×)

Treatment with candlenut oil revealed a dose-dependent histological improvement. Group C (0.5 mL/200 g BW) showed moderate mucosal preservation with partial epithelial restoration. Group D (1.0 mL/200 g BW) exhibited improved epithelial continuity and reduced inflammatory cell infiltration, while Group E (2.0 mL/200 g BW) demonstrated near-complete mucosal integrity and minimal inflammation. Although not equivalent to Group A, the histological appearance of Group E was notably better than that of Group B.

These histological findings align with prior research on flavonoid-rich plant oils such as *Nigella sativa* and *Camellia sinensis*, which demonstrated mucosal preservation and reduced gastric lesion formation in NSAID-induced models(29,30). Likewise, flavonoids in *Citrus aurantium*—including hesperidin and quercetin—have been shown to mitigate oxidative and inflammatory damage in gastric tissues(31,32).

### Quantitative Evaluation of Mucosal Injury

Histopathological scoring using Roger’s criteria (Table 1) corroborated the visual findings. The negative control group (Group B) exhibited the highest mean damage score (5.40 ± 1.673), significantly higher than the healthy control (2.20 ± 0.837, p < 0.01). Groups C, D, and E exhibited progressive reductions in injury scores (5.00 ± 0.707, 4.20 ± 0.837, and 4.00 ± 1.000, respectively). **One-way ANOVA** confirmed a significant overall group difference (p = 0.001).

Post hoc Tukey HSD test revealed a significant difference between Groups B and A (p < 0.01), validating aspirin’s mucosal toxicity. No significant differences were found between Group B and the treatment groups (p > 0.05); however, Group E differed non-significantly from Group A (p = 0.096), suggesting partial mucosal restoration.

The biological trend toward dose-dependent protection is consistent with prior studies using *Nigella sativa* or *Hibiscus cannabinus* (kenaf) seed oil, where escalating doses were associated with reduced ulceration indices(33,34). This trend likely reflects improved tissue bioavailability of bioactive constituents—such as polyphenols and unsaturated fatty acids—at higher doses.

### Mechanistic Insights and Implications

The observed histological restoration is plausibly mediated by anti-inflammatory and antioxidant effects of bioactive compounds in candlenut oil. Oils rich in linoleic and oleic acids—such as safflower and perilla oils—are known to attenuate NSAID-induced injury by reducing malondialdehyde (MDA) levels and enhancing superoxide dismutase (SOD) and glutathione peroxidase (GPx) activity (35,36). These bioactive pathways correspond with the presumed activity of *A. moluccanus*, which has been reported to contain polyunsaturated fatty acids, flavonoids, and tocopherols(37–39).

Despite the lack of statistical significance in pairwise comparisons, the consistent histological improvement suggests a pharmacologically relevant effect. Such patterns are not uncommon in NSAID-induced ulcer models where protective trends may emerge visually despite small sample sizes(40–42). The downward trajectory in histological scores across treatment groups supports the hypothesis that candlenut oil exerts cytoprotective effects through modulation of oxidative and inflammatory pathways(43).

Future studies aiming to validate mechanisms in gastroprotective research should indeed focus on quantifying biomarkers such as malondialdehyde (MDA), tumor necrosis factor-alpha (TNF-α), interleukin-1 beta (IL-1β), and cyclooxygenase-2 (COX-2) expression. MDA serves as a marker for oxidative stress, which is closely linked to inflammation and various chronic diseases, including gastric cancer (44,45).

### Safety Considerations and Translational Potential

Processed candlenut oil has demonstrated a favorable preclinical safety profile. Toxicological studies in rodents show no adverse effects on lipid profiles or hepatic biomarkers at moderate doses(45,46). However, phytochemical standardization remains essential, particularly due to mild toxicity reported with raw or unrefined seeds. Ensuring consistent levels of linoleic acid, oleic acid, and total flavonoids would improve reproducibility and satisfy regulatory expectations for botanical therapeutics(47–50).

The accessibility and affordability of *A. moluccanus* oil, especially in tropical regions where synthetic gastroprotectants are limited, enhance its value as a complementary therapy(17). Further development should focus on formulation optimization (e.g., softgel capsules or microencapsulation), dose-escalation studies, and long-term toxicity assessments to enable clinical translation.

### Limitations

This study has several limitations. Methodological limitations include the lack of phytochemical standardization of candlenut oil, absence of systematic documentation of gastric mucosal morphology, and omission of a positive control group (e.g., omeprazole or sucralfate) to contextualize gastroprotective effects. Only male rats were used, preventing evaluation of sex-based differences, and relatively high intra-group variability may have reduced statistical power. Histological assessment, although performed by two independent blinded observers including a board-certified pathologist, did not include formal inter-observer reliability testing.

Mechanistic limitations include the absence of quantitative assessment of inflammatory cytokines, oxidative stress markers (e.g., TNF-α, IL-1β, MDA), gastric pH, and mucus thickness, which restricts mechanistic interpretation of the observed gastroprotective effects.

Future studies should incorporate positive controls, both sexes of animals, larger sample sizes with power analysis, standardized phytochemical characterization, systematic morphological documentation, and relevant biomarker analyses to validate mechanisms and enhance translational relevance.

## CONCLUSION

The findings of this study indicate that *Aleurites moluccanus* oil may confer protective effects against aspirin-induced gastric mucosal injury in rats, as reflected by histological improvements in mucosal integrity and inflammatory cell infiltration. Although not all results reached statistical significance, the dose-dependent trend suggests potential pharmacological relevance. These preliminary results support further investigation into the underlying mechanisms and therapeutic value of *A. moluccanus* oil as a gastroprotective agent. In addition to further mechanistic studies, acute, sub-acute, and chronic toxicity evaluations following OECD guidelines are planned to establish a comprehensive safety profile before advancing candlenut oil toward clinical evaluation.

## ACKNOWLEDGMENTS

The authors express their sincere gratitude to the Pharmacology Laboratory, Faculty of Medicine, Universitas Jenderal Soedirman, for providing technical support and research facilities. Appreciation is also extended to the Medical Research Ethics Committee of Universitas Jenderal Soedirman for ethical clearance (Approval No. 002/KEPK/PE/XI/2022). The authors acknowledge the contributions of laboratory technicians and support staff involved in animal handling and histopathological preparations during the study.

## AUTHOR CONTRIBUTIONS

Eman Sutrisna conceptualized the study, designed the experimental protocol, and prepared the initial manuscript draft. Mohammad Rizky Ariesto and Hidayat Sulistyo conducted the laboratory experiments, animal handling, and histopathological procedures. Iwan Purnawan supervised the overall research process, performed data analysis, and critically revised the manuscript. All authors reviewed and approved the final version and agree to be accountable for all aspects of the work..

## ETHICAL APPROVAL

This study was conducted under ethical clearance issued by the Medical Research Ethics Committee, Faculty of Medicine, Universitas Jenderal Soedirman, with approval number 002/KEPK/PE/XI/2022. All animal procedures complied with national guidelines for the care and use of laboratory animals.

## DATA AVAILABILITY

All data generated and analyzed during this study are included in this published article. Further details can be provided by the corresponding author upon reasonable request.

## PUBLISHER’S NOTE

All claims expressed in this article are solely those of the authors and do not necessarily represent those of the publisher, the editors, or the reviewers. The publisher remains neutral with regard to jurisdictional claims in published institutional affiliation.

## USE OF ARTIFICIAL INTELLIGENCE (AI)-ASSISTED TECHNOLOGY

Artificial intelligence (AI) tools were utilized solely to improve the clarity, coherence, and logical structure of the manuscript’s language. The use of AI did not involve generation or fabrication of data, results, or scientific content. All research findings, analyses, and interpretations presented in this article are entirely based on original experimental work conducted by the authors.

